# Spargel/dPGC-1 influences cell growth through the E2F1-mediated endocycle pathway

**DOI:** 10.1101/2025.08.02.668293

**Authors:** Md Shah Jalal, Atanu Duttaroy

## Abstract

The endocycle is a specialized variant of the eukaryotic cell cycle program observed in various tissues across diverse organisms ranging from insects to mammals. The endocycle promotes cellular growth by alternating between the synthesis (S) and gap (G) phases, completely bypassing mitosis (M phase). E2F1 serves as a master regulator of the endocycle in *Drosophila* salivary glands, whereas the TOR signaling pathway controls the levels of the E2F1 protein post-transcriptionally. Cellular growth in tissues that undergo the endocycle is also dependent on nutrient availability. *Drosophila* Spargel (dPGC-1) is orthologous to a group of transcriptional co-activators in vertebrates known as PGC-1. In flies, Spargel influences cell growth through the Insulin signaling pathway via TOR. However, the mechanisms by which Spargel regulates endocycle-mediated growth have yet to be established. Here, we report an essential role of Spargel in the *Drosophila* larval salivary gland, which influences E2F1-mediated cellular growth. To elucidate the role of Spargel in the salivary gland, we performed FLP/FRT-mediated clonal analysis and found that a cell-specific loss of Spargel leads to smaller nuclei with reduced DNA content due to the early termination of DNA replication. Further, the selective absence of Spargel abrogates the expression of a key DNA replication factor called E2F1, which promotes G1→S transition in endocycle. Thus, mechanistically, Spargel plays a key role in cell growth by positively influencing the endocycle process through the E2F1 pathway.

## Introduction

As growth occurs during development, the size and number of cells ultimately determine the body size of an adult animal (Tennessen and Thummel, 2011; Conlon and Raff, 1999). *Drosophila* is a recognized model for studying growth because its juvenile larval stage undergoes dramatic changes, leading to a remarkable ∼200-fold increase in body mass (Tennessen and Thummel, 2011; Church & Robertson, 1966). The initial growth of *Drosophila* embryos is mitotic, while post-embryonic growth relies entirely on the increase in cell size of larval endoreplicating tissues (Gupta et al., 2013; Church and Robertson, 1966). *Drosophila* larval tissues shift from mitosis to a different cell cycle program called the endocycle, which includes only the S and G phases and omits cytokinesis. (Edgar et al., 2014).

*Drosophila* larvae possess a pair of salivary glands, each comprising approximately 100 cells (Andrew et al., 2000). Once cells are committed to forming the salivary glands, no cell division or cell death occurs within these glands (Andrew et al., 2000). Throughout larval development, the salivary gland cells increase in size through endocycle (Loganathan et al., 2021; Shu et al., 2018). The endocycle leads to polyploid cells with multiple copies of their genome, thereby driving growth (Shu et al., 2018). The transcription factor E2F1 plays a central role in regulating the endocycle in *Drosophila* larval salivary gland (Kim et al., 2021), and the three upstream effectors of E2F1, namely Myc, the target of rapamycin (TOR), and receptor tyrosine kinases (RTK), regulate the endoreplication pace (Edgar et al., 2014).

The PGC-1 group of transcriptional coactivators serves as a master regulator of mitochondrial biogenesis and energy metabolism in mammals (Lin et al., 2003; Lin et al., 2002; Andersson and Scarpulla, 2001). The discovery of an ancestral PGC-1, Spargel/dPGC-1 (Tiefenböck et al., 2010), led us to explore the biology of this essential transcription co-activator in *Drosophila* by harnessing the power of fly genetics and the widely used RNAi technique. Previous studies have documented the indispensable role of Spargel/dPGC-1 in germ cell growth and its significance for female fertility (Basar et al., 2019) as well as in eggshell biogenesis (Jalal and Duttaroy, 2024). In this report, we highlight that Spargel/dPGC-1, located in the cytoplasmic compartment, plays a positive role in the endocycle process of larval tissues during the postembryonic development of *Drosophila*. Thus, Spargel is involved in cellular and, consequently, organismal growth through endocycle.

## Results

### Spargel expression pattern in developing somatic tissues like muscle, gut, and salivary gland

Late-stage embryonic lethality of *spargel*(*srl*) null embryos specifies a zygotic role for this transcriptional co-regulator during *Drosophila* development (Roy et al., 2023). Earlier studies documented that Spargel is highly expressed in the nuclei of ovarian germ cells and follicle cells in adult females (Basar et al., 2019). In contrast, the expression of Spargel in the cytoplasm of fat body cells has also been reported (Tiefenböck et al., 2010). Using Spargel monoclonal antibody (Basar et al., 2019), we performed a detailed survey of Spargel expression in major larval tissues, including salivary gland, gut, and muscles (**Fig. 1A-C)**. The presence of Spargel in the cytoplasm of these larval tissues supports previous findings that Spargel is localized within cytoplasmic compartments in all somatic cell tissues. However, more dynamic spatiotemporal changes in cytoplasmic Spargel expression pattern were observed in the larval salivary gland during larval to pupal transition. In wandering 3^rd^ instar larvae, Spargel is expressed predominantly in the proximal region of the glands (**Fig. 1D**). No Spargel expression is evident in the distal region of the gland during this time (**Fig. 1D**). However, in pre-pupal and pupal stages, the cytoplasmic expression pattern of Spargel expands from the proximal to the distal region, and eventually it expresses throughout the glands (**Fig. 1E-F**).

**Figure 1:**
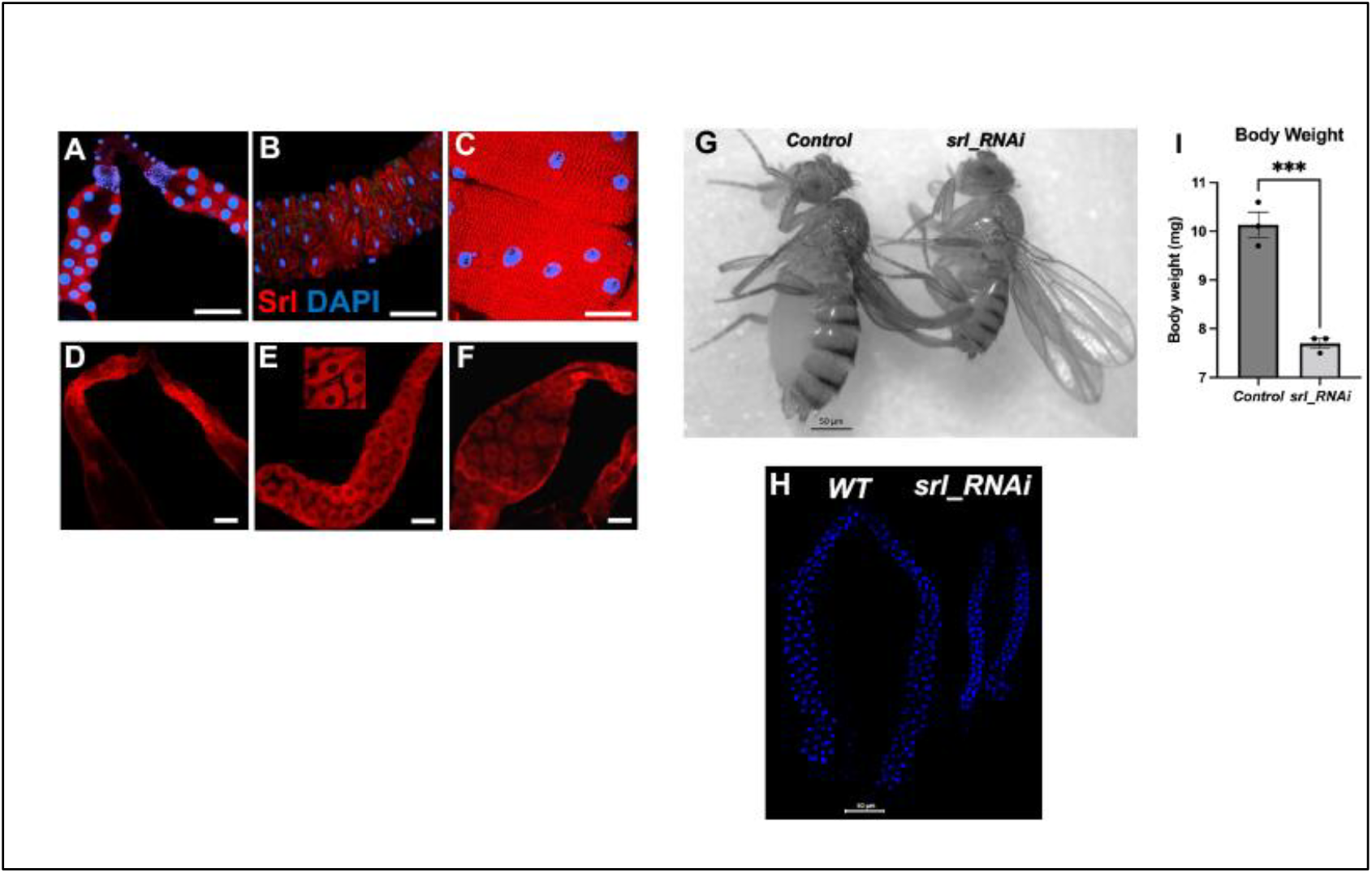
Spargel protein expression profile in larval somatic tissues. Immunostaining with the Spargel-specific monoclonal antibody (7A10) reveals cytoplasmic localization of Spargel (red) in all examined larval somatic tissues: (A) salivary gland, (B) midgut, and (C) muscle tissues from w1118 wandering 3rd instar larvae. Nuclei were counterstained with DAPI (blue). Scale bars are 100 µm for panels A and B, and 50 µm for panel C. (D-F) The distribution and expression pattern of Spargel in the salivary gland during later stages of larval development. In 3rd instar larvae, Spargel expression appears in the proximal region of the glands (D). In contrast, during prepupal and pupal stages, a notable extension of cytoplasmic Spargel expression spreads from the proximal to the distal region of the gland (E), persisting until gland histolysis (F). The scale bar is 100 µm. (G-H) Ubiquitous (G) and salivary gland-specific (H) knockdown of spargel causes reduced body size and smaller, thinner glands due to stunted growth. A comparison of the average body weight between control and srl-RNAi flies confirms a significant reduction in body weight.

### Ubiquitous and tissue-specific loss of Spargel displays a growth defect

To investigate what role cytoplasmic Spargel is playing in the developing larval tissues, ubiquitous and salivary gland-specific knockdown of *spargel* was performed with *tubulin_gal4* and *AB1_gal4* drivers, respectively. In each case, *spargel* knockdown results in somatic growth arrest. The global knockdown of *spargel* produces smaller body size and significantly underweight flies (**Fig. 1G, I**), whereas the salivary gland-specific knockdown of *spargel* results in rudimentary glands (**Fig. 1H**) that mimic the *spargel* knockdown phenotype in ovaries (Basar et al., 2019). Therefore, the global and tissue-specific suppression of *spargel* indicates a correlation between *spargel* and growth, as the downregulation of *spargel* leads to reduced growth across all examined tissues. These observations raise the question: Is Spargel fundamentally associated with growth? To address the question, we investigated the cellular requirements of Spargel using a widely employed mitotic clonal technique.

### Cell-specific loss of *spargel* supports its role in endocycle-mediated cell growth

The mitotic clonal technique (Golic and Lindquist, 1989) is used to elucidate the requirement of Spargel in the salivary gland cells and other somatic tissues by extension. Loss of GFP reporter and Spargel signal (immunostained with Spargel-specific antibody) confirmed the absence of Spargel (*srl*^*null*^) in specific salivary gland cells **(Fig. 2A-A**’ **and B-B**’). In contrast, the adjacent salivary gland cell with wild-type Spargel expression serves as a control **(Fig. 2A-A**’ **and B-B**’). *srl*^*null*^ cells in the salivary gland cells have nuclei that are significantly smaller in size compared to the adjacent control cells (**Fig. 2C-C**’). Quantification of nuclear DNA content as measured through DAPI intensity of GFP negative nuclei (**Fig. 2D-D**’) established that *srl*^*null*^ clones carry half the DNA content compared to the adjacent control cells (**Fig. 2E)**. Salivary gland cells are polyploid. The reduced DAPI intensity and smaller nuclear size of the *srl*^*null*^ cell clones prompted us to investigate the endoreplication status of these cells.

**Figure 2:**
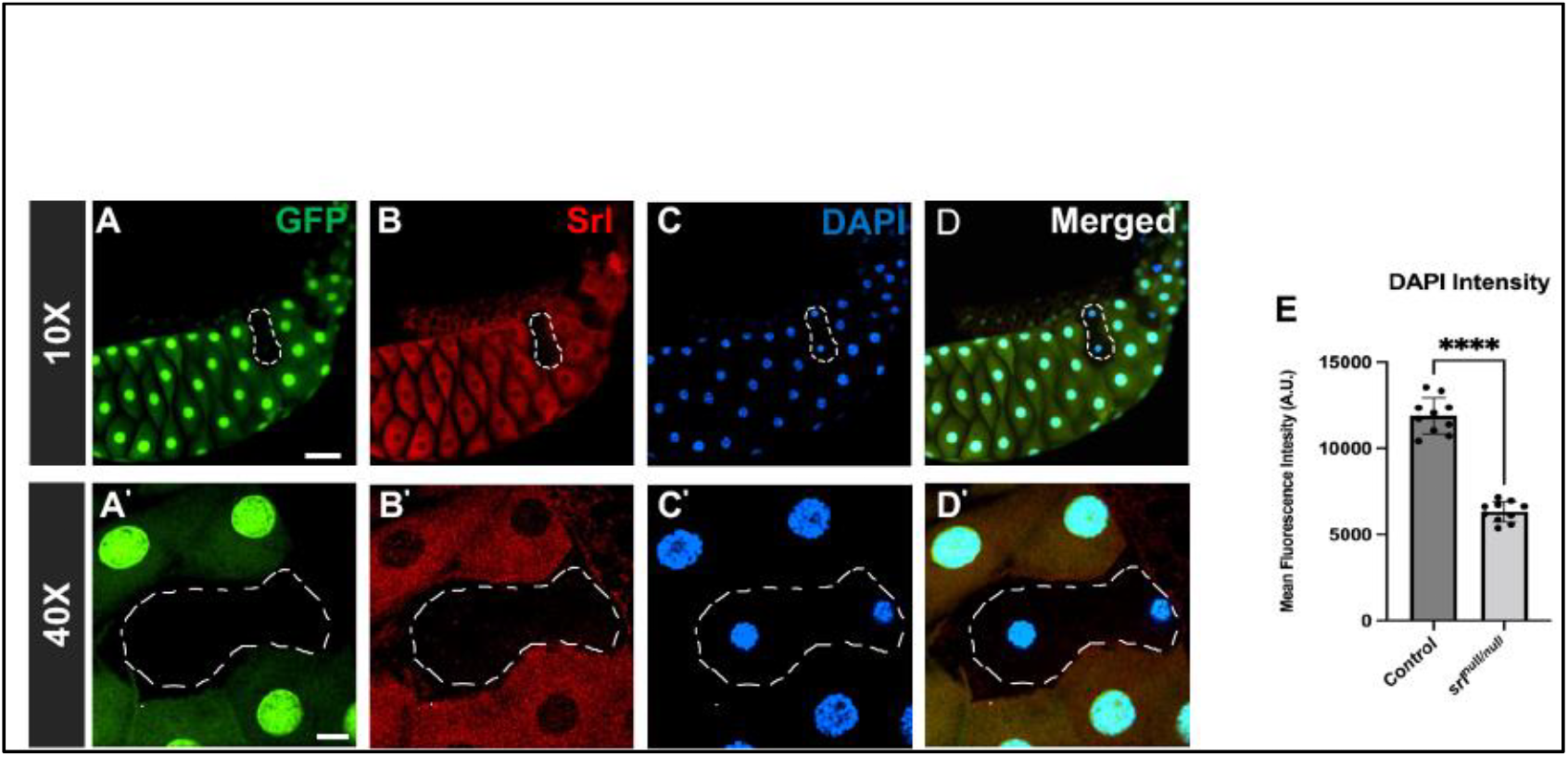
Cell-autonomous loss of *spargel* function in the salivary gland leads to smaller nuclear size. Panels A-D and A’-D’ display the same tissue at 10x and 40x magnification, respectively, showing that nuclear size reduction is evident at both tissue-wide and cellular levels. (A-A’) *srl*^*null*^ clones are GFP negative, whereas GFP-positive cells are *control* cells. (B-B’) GFP negative cells are also negative for Spargel expression (red), (C-C’) DAPI positive (blue) *srl*^*null*^ nuclei are much smaller in size compared to the adjacent control cell nuclei. (D-D’) GFP-negative and DAPI-positive nuclei have smaller nuclei. (E) Quantification of DAPI intensities in the nuclei of *control* cells compared to the *srl*^*null*^ clone cells. The number of dots represents the number of nuclei counted in each group. The error bars represent SEM, and the P value is < 0.0001. n=10. White lines across all panels indicate the clone boundary. Scale bar, 100µm (top panel) and 20µm (bottom panel).

The endoreplication status of *srl*^*null*^ cells was tested through EdU incorporation, which labels the S-phase nuclei (Buck et al., 2008). EdU incorporation is evident in control nuclei, indicating that at least one round of DNA replication has happened during the labeling period (**Fig. 3A-D**). In contrast, the total absence of EdU incorporation in *srl*^*null*^ cell clones validates the reduced endoreplication status of these cells (**Fig. 3B**). Quantifying the intensity of EdU-positive and negative nuclei in both control and clonal groups **(Fig. 3E)** shows that the absence of Spargel almost completely halts DNA replication in the salivary gland.

**Figure 3:**
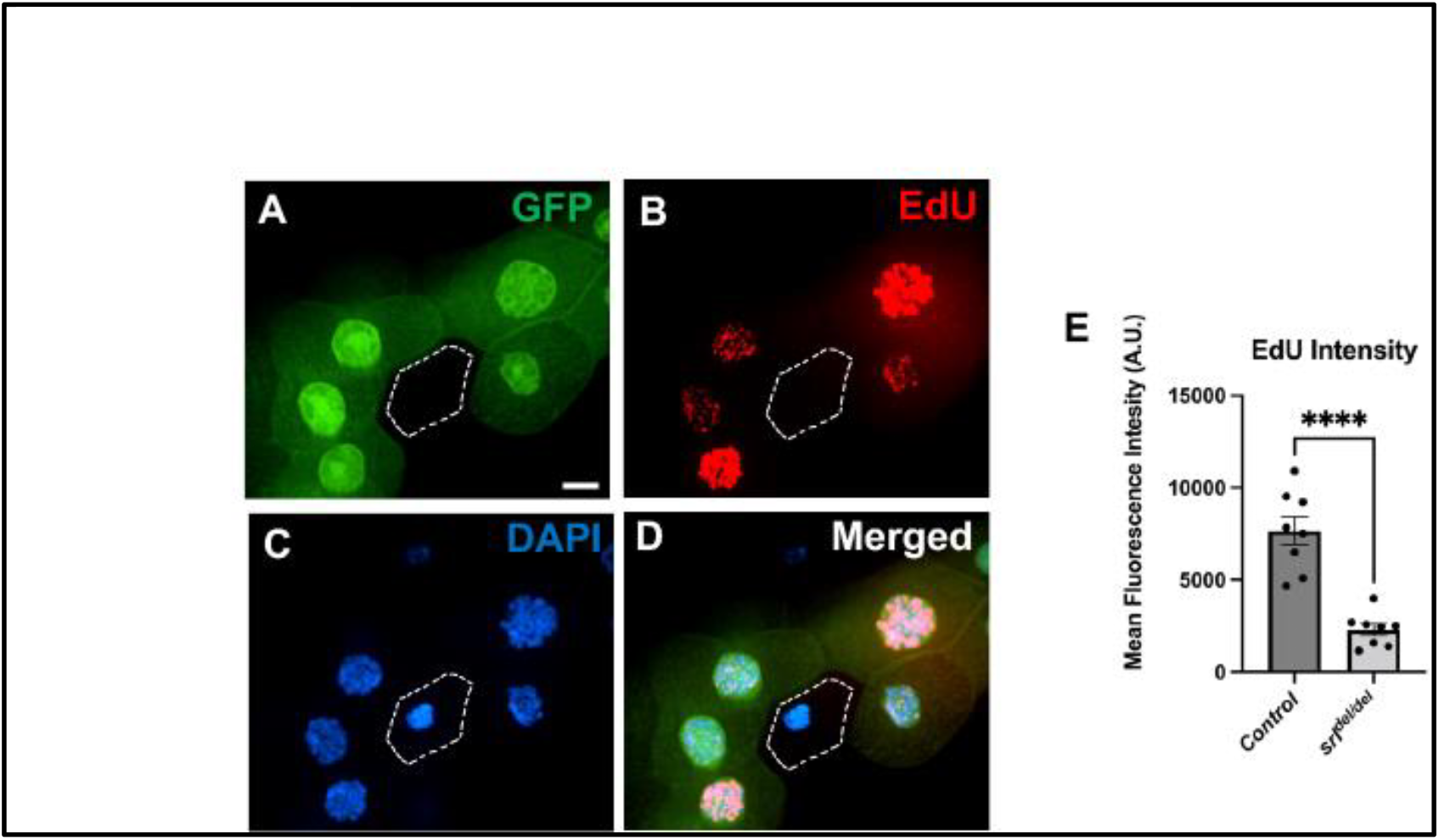
*srl* ^*null*^ nuclei in the salivary gland undergo reduced endoreplication. EdU incorporation into the clonal salivary gland to detect S-phase nuclei. (A) GFP-negative cells in the salivary gland represent the *srl*^*null*^ clone. (B) Images of salivary glands following EdU incorporation (red) support active EdU incorporation in *control* nuclei (GFP positive). However, *srl*^*null*^ clones are lagging in their endoreplication status, as evident from very little to no EdU incorporation in these cells (white marked region). (C) Representative images stained with DAPI show a smaller nucleus in the *srl*^*null*^ clone, as indicated by the lack of GFP. (D) Merged images. (E) The comparison of EdU intensity in the *control* cells to *srl*^*null*^ clone cells. The number of dots represents the number of nuclei. The error bars represent SEM, and the P value is < 0.0001. n=8. White lines across all panels indicate the clone boundary. Scale bar 20µm.

### E2F1, a core regulator of endocycle, is nearly abolished in *srl*^*null*^ cell clones

E2F1 is a core regulator of endocycle in the salivary gland, as downregulation of E2F1 results in reduced endocycle (Zielke et al., 2013). Since cell-specific loss of function of *spargel* causes reduced DNA content by negatively influencing DNA replication, this prompted us to profile E2F1 expression in the *srl*^*null*^ cell clones. E2F1 expression is abrogated in *srl*^*null*^ cell clones, while adjacent wild-type cells exhibit normal E2F1 expression **(Fig. 4A-C)**. In addition, measurements of E2F1 intensity indicate that E2F1 proteins are nearly undetectable in the *srl*^*null*^ cell clones. This data is quite significant, which leads us to conclude that Spargel is upstream of E2F1. Furthermore, Spargel and E2F1 work together to maintain proper endoreplication status and regulate cell growth.

**Figure 4:**
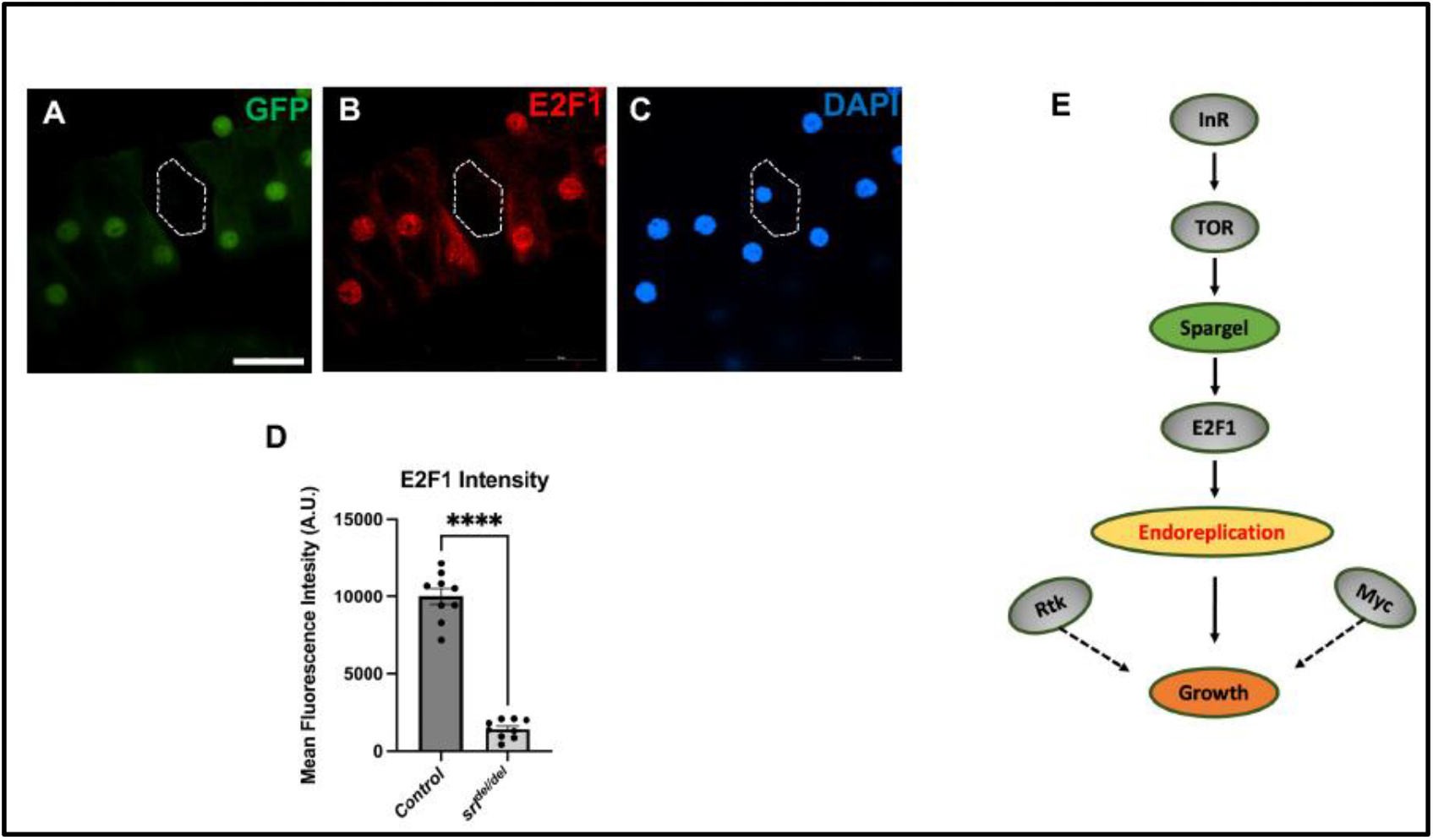
E2F1 expression is nearly abolished in *srl*^*null*^ cells. (A-C) Immunostaining with E2F1 antibody in the salivary gland containing the *srl*^*null*^ clone. Absence of GFP expression, marked with a white dotted line, indicates the *srl*^*null*^ clone (A). E2F1(red) and DAPI-stained (blue) nuclei show a smaller nucleus in the *srl*^*null*^ clone, as indicated by the lack of GFP expression in the nuclei of the *control* and the *srl*^*null*^ clone (B and C, respectively). (D) E2F1 intensity in the *control* cells compared to the *srl*^*null*^ clone cells. The number of dots represents the number of nuclei. The error bars represent SEM, and the P value is < 0.0001. n=9. White lines across all panels indicate the clone boundary. Scale bar 100µm. (E) Proposed mechanism of Spargel action in the endocycle-mediated pathway in the salivary gland. Spargel, being upstream, influences the expression of E2F1, which in turn positively influences the endoreplication process.

## Discussion

Endocycle is an alternative cell cycle program that allows cells to increase in size by repeatedly replicating their DNA content without undergoing cellular division (Edgar et al., 2014). Endocycle typically consists of only S (synthesis) and G (gap) phases, which results in polyploid cells with multiple copies of their genome (Shu et al., 2018). Skin, placenta, liver, and blood cells undergo endoreplication in mammals during development (Zielke et al., 2013). Endocycle is essential for determining adult body size in nematodes, which is also common in arthropods, including insects, and is well characterized in *Drosophila* (Zielke et al., 2013). Endocycle and polyploidy are essential in many *Drosophila* larval tissues, including the salivary gland, gut, fat body, and adult tissues like ovarian nurse and follicle cells, gut, and glia for achieving normal sizes (Øvrebø et al., 2022; Kim et al., 2021; Orr-Weaver, 2015; Zielke et al., 2013; Fox & Duronio, 2013; Zielke et al., 2011).

Endocycle and the mitotic cycle share common regulators like the transcription factor E2 promoter binding factor (E2F) and cyclin-dependent kinases (CDKs) (Lee et al., 2009; Lilly & Duronio, 2005). The *Drosophila* salivary gland serves as a unique tissue for studying the endocycle, where it has been determined that the transcription factor E2F1 plays a central role in regulating this process (Kim et al., 2021). The salivary gland cells lacking *E2F1* block endoreplication, and overexpression of *E2F1* results in increased endoreplication, leading to the formation of hyperpolyploid cells (Zielke et al., 2011). In this observation, we established that E2F1 expression is Spargel dependent, meaning E2F1 functions downstream of Spargel **(Fig. 4E)**. Since E2F1 plays a central role in endocycle of nuclear DNA and ultimately in cellular and tissue growth, E2F1 being downstream of *spargel* easily explains the reduced cell, tissue, and eventually organismal growth resulting from the lack of Spargel protein.

The endoreplication pace in the *Drosophila* salivary gland is indirectly controlled by three upstream effectors: Myc, the target of rapamycin (TOR), and receptor tyrosine kinases (RTK) (Edgar et al., 2014), as DNA replication is affected by each. These upstream factors also regulate the translation rate of *E2F1*, and alterations in these effectors lead to endocycle anomalies (Edgar et al., 2014). In a key observation, Tiefenböck et al. (2010) claimed that in fat body cells, growth is enhanced by overproduction of Insulin Receptor (InR), whereas the significant cell growth resulting from InR overexpression is abrogated in *spargel* hypomorph (*srl*^*1*^). Simultaneously, gene array analysis supported the induction of *srl* transcript levels in InR-overexpressing cells. In addition to InR, Spargel gain-of-function rescued the cell size and growth defects of other members in the insulin signaling pathway, such as TOR and S6K, in a cell-autonomous manner (Mukherjee et al., 2013). All this evidence points to the direct role of Spargel downstream of the Insulin-TOR signaling pathway (Tiefenböck et al., 2010; Mukherjee et al., 2013). Thus, like other members of the insulin signaling pathway, Spargel can also control cell growth. But the significant question that remains is how? Exclusion of E2F1 from *srl*^*null*^ cell clones provides the strongest evidence that Spargel is involved in E2F1-mediated endoreplication and thus controls cell growth. Future work will reveal: (1) how a transcription co-activator (Spargel) and transcription factor (E2F1) work together to control cellular growth, and (2) how cytoplasmic Spargel can control the action of a transcription factor located in the nuclear compartment.

## Materials and methods

### Drosophila strains and genetics

All the fly stocks were raised on standard cornmeal agar medium at a constant temperature (23°C). The following Drosophila melanogaster stocks were obtained from BDSC: *w*^*1118*^ *(w[1118]), y[1]; P{y[+mDint2] w[BR*.*E*.*BR]=SUPor-P}srl[KG08646] ry[506]/TM3, Sb[1] Ser[1]* (BDSC #14965), *w[*]; P{w[+mC]=matalpha4-GAL-VP16}V37* (BDSC #7063), w[1118]; P{w[+mC]=UASp-EGFP.srl}/CyO (Duttaroy lab), *y[1] sc[*] v[1] sev[21]; P{y[+t7*.*7] v[+t1*.*8]=TRiP*.*HMS00857}attP2* (BDSC #33914).

### Generation of *srl*^*null*^ cell clones

*spargel* loss-of-function (*srl*^*null*^) clones were generated in the larval tissues using mitotic recombination. A CRISPR-mediated new FRT insertion at FRT81F6 was generated first because the canonical FRT82B6 insertion is near the *spargel* locus, which impedes mitotic recombination (Roy et al., 2023). *ry,hsflp,y,w*;;FRT81F6,srl*^*del*^ */TM3,ry, Sb* males were crossed to *w*; P{w[+mC]=Act5C-GAL4}25FO1/CyO; FRT81F6, UAS-GFP,w+/TM3, Sb* virgin females. The 4-5 hours F1 embryos from the above cross were collected on grape agar plates and heat-shocked once for 1.5 hours at 37°C. Heat-shocked samples were raised in an incubator (23°C) for larval growth. Then, the larval tissues, including the salivary glands, were dissected from the desired larval stage (wandering 3^rd^ instar) for further analysis. *srl*^*null/null*^ clones in the salivary gland cells were identified by the absence of a GFP marker.

### Immunostaining and antibodies

The salivary glands were dissected in Grace’s Insect Medium (Life Technologies, cat. #11605-094), fixed in 1X PBS with 4% paraformaldehyde (Ted Pella, cat. #18505) for 20 minutes at room temperature (note: for anti-dE2F1, tissues were fixed (30min), washed and blocked at 4^0^C). Tissues were then washed three times in 1XPBS with 0.3% Triton-X-100 (PBST) and blocked in a 1% BSA in PBST (PBTB) for 1 hour at room temperature. The tissues were incubated in PBTB with an appropriate primary antibody overnight at 4 ^°^C, then washed three times in PBST at room temperature. The secondary antibody incubation was carried out for 1 hour at room temperature in the dark. Then, the tissues were washed three times in PBTB and mounted in VECTASHIELD antifade mounting medium with DAPI (Vector Laboratories, cat. #H-1200-10) for imaging. The following primary and secondary antibodies were used in this study: mouse anti-Spargel (7A10), rabbit anti-E2F1 (Maxim Frolov lab), rhodamine Red™-X goat anti-mouse (Molecular Probes, Life Technologies), rhodamine Red™-X goat anti-rabbit (Molecular Probes, Life Technologies).

### EdU incorporation

A Click-iT EdU Alexa Fluor 488 Imaging Kit (Thermo Fisher Scientific, cat. #C10337) was used to incorporate 5-ethynyl-2’-deoxyuridine (EdU) into the *Drosophila* salivary gland following the manufacturer’s instructions.

### Microscopy

The immunostained and EdU-labeled samples were imaged with a Nikon Ti-E-PFS inverted confocal microscope equipped with a Yokogawa CSU-X1 spinning disk using the 40x (oil), 20x, and 10x Plan Apo λ objective lenses. The NIS Elements Ar imaging software was used to adjust the gamma, brightness, and contrast. Adult flies and the DAPI-stained salivary glands were photographed and acquired with a Zeiss Axioskop 2 plus microscope in brightfield mode, equipped with a PhotoFluor LM-75 light source using a 5x or 10x Zeiss Fluar objective lens.

### Statistical analysis

Quantification and statistical evaluation were accompanied by at least three independent replicates collected under identical settings. Students’ t-tests were used to determine statistical significance and compare the two data groups. Statistical software, including GraphPad Prism 9 and Microsoft Excel, was used to analyze and generate bar charts. The following p-values were evaluated for statistically significant differences: * P ≤ 0.05 ** P ≤ 0.01 *** P ≤ 0.001 **** P ≤ 0.0001.

